# Upstream ribosome impediments activate roles of internal Shine-Dalgarno sequence for translation initiation in *E. coli*

**DOI:** 10.1101/2023.07.05.547755

**Authors:** Xin Li, Huihui Wang, Yan Chen, Yanyan Zhang, Jie Liu, Dan Zhou, Zhihua Wu, Meihao Sun

## Abstract

The initiation of protein translation, one of the four phases of translation, had been proved to be the rate limiting step for translation. The specific interaction of bacterial 30S ribosomal subunit with Shine-Dalgarno sequence (SD) contributes the initiation significantly. It had been shown that SD-like sequence in coding region of mRNA, designated as internal SD, can direct translation initiation as regular SD does. The ribosome impediments were demonstrated to be one of the factors contributing the non-uniform translation elongation rate, but their effects on internal SD translation initiation role (ISTIR) remains unclear. To investigate effects of upstream ribosome impediments on ISTIR, a fragment consisting of pyruvate kinase (PK) gene, translation initiation needed mRNA elements (including an A/U-rich region, SD sequence and start codon AUG) and red fluorescent protein (RFP) gene was constructed and RFP expression levels representing translation initiation efficiency induced by internal SD was analyzed. Surprisingly, RFP expression was not detected with this primary construct, further experiments of inclusion of stem loop structure preceding to internal SD or co-expression of engineered RNA binding scaffold (ERBS) targeting to preceding sequence of internal SD could activate ISTIR. These results suggest that upstream ribosome impediments would activate ISTIR to initiate the downstream gene translation, which manifests the potential for developing new method to test interactions between RNA binding proteins and their target RNA molecules *in vivo*.

## Introduction

Protein translation in prokaryotes could be divided into four phase: initiation, elongation, termination and ribosome cycling (Rodnina 2018). For the canonical translation initiation, interaction between ribosomal protein S1 and A/U-rich specific regions of mRNA is followed by 30S ribosomal subunit scanning of mRNA. Once ribosome meets another the specific purine-rich region on mRNA, called ribosome binding site and referred to as Shine-Dalgarno sequence (SD) (Shine and Dalgarno 1974), locating 6-8 bases upstream of the start codon, SD would interact with the CU rich anti-Shine Dalgarno sequence at the 3′ tail of 16S rRNA (Chen et al. 1994; Laursen et al. 2005). Aligning the start codon to the P site of the 30S subunit, recruiting initiation factors and 50S ribosomal subunit lead to formation of the initiation complex (Wei et al. 2017) and finish the translation rate limit step, the translation initiation.

Elongation process includes steps of aminoacyl tRNA binding, peptide bond formation and translocation, resulting incorporation of one amino acids into the nascent peptide. The process of elongation occurs at a non-uniform rate and that the speed of elongation could be regulated by codon choice and distribution (Mohammad et al. 2016), mRNA secondary structure (Wen et al. 2008), composition of the aa-tRNA pool (Rudorf et al. 2014; Vieira et al. 2016; Dykeman 2020) and amino acid composition of nascent peptide (Ito and Chiba 2013; Wilson et al. 2016; Riba et al. 2019). Specific sequences, such as SD-like sequence in coding region of mRNA (internal SD), causing rRNA– mRNA hybridization had been proposed for regulating the translation elongation process (Ponnala 2010). Internal SD had been documented to arrest the translation by binding and pausing ribosome at the specific site, decrease the likelihood of ribosome drop off and frame shift the translation (Weiss et al. 1988; Larsen et al. 1994; Wen et al. 2008; Ponnala 2010; Li et al. 2012). Not only that internal SD was also proposed to initiate translation of downstream gene, based on the distance between internal SD and the start codon, and the fact of high frequency of C-terminal internal SD in upstream gene succeeded by overlapping downstream genes (Diwan and Agashe 2016), but also it was found to initiate protein translation in *E. coli* (Cicek et al. 2013; Leith et al. 2019). The ribosome impediments, such as strong secondary structures (Wen et al. 2008), were demonstrated to be one of the factors contributing the non-uniform translation elongation rate, but their effects on internal SD translation initiation role (ISTIR) remains unclear.

In this study, ribosome impediments of stem loop secondary structure and RNA binding protein were used to investigate their effects on ISTIR with *E. coli* expression system. The testing pET vector was constructed by incorporation of internal SD between genes of pyruvate kinase (PK) and red fluorescent protein (RFP), and ISTIR was analyzed by checking RFP expression after IPTG induction. The results surprisingly showed that no RFP signal was detected after IPTG induction, which indicates the absence of ISTIR for the constructed internal SD. But introduction of stem loop structure in mRNA or co-expression of RNA binding protein which binds to the upstream of internal SD would induce the RFP expression, a signal indicating the activation of ISTIR. These results clearly demonstrated that internal SD presence is not the sole factor necessary to induce downstream mRNA expression, and the translation impediment located upstream of internal SD would activate ITIR.

## Materials and methods

### Materials

Lysozyme, 1,4-Dithiothreitol, pepstatin A, isopropyl β-D-1-thiogalactopyranoside (IPTG), and phenylmethanesulfonyl fluoride (PMSF) were obtained from Amresco (USA). KCl, plasmid preps kit and bacterial genomic DNA isolation kit were acquired from Shanghai Sangon (Shanghai, China). Oligoes and coding sequence of engineered RNA binding scaffold1 (ERBS1) were synthesized by Shanghai Sangon. DNA restriction enzymes, DNA markers, T4 DNA ligase and polymerase were from Takara (Shiga, Japan). Antibiotics were purchased from Oxford LTD (Hampshire, England). Bacterial strains and pETDuet-1 were purchased from Novagen (New Jersey, USA). All other chemicals were of analytical grade and obtained from domestic companies.

### Construction of expression vectors

*E. coli* genomic DNA was purified with bacterial genomic DNA isolation kit following the manual instruction. PK coding sequence was amplified by PCR with *E. coli* genomic DNA as the template and the primers of PK-F and PK-R (suppl table1). The amplified fragment was inserted into the second polyclonal site of pETDuet-1 after *Nde* I and *Kpn* I digestion and further ligated by T4 ligase, resulting pETDuet-1-PK expression vector.

RFP coding sequence and internal SD was amplified by two rounds PCR with His9-RFP plasmid (Zhang et al. 2015) as template and primers of RFP-F2 and RFP-R for the first round. The second round PCR was made with first round PCR product as template and primers of RFP-F1 and RFP-R. Product of second round PCR was incorporated into vector pETDuet-1-PK after *Kpn* I and *Xho* I digestion and further ligated by T4 ligase, resulting pETDuet-1-PKRFP expression vector (suppl data1).

Introduction of *Eco*R I restriction site into pETDuet-1-PKRFP expression vector was made with primers ER1 and ER2, yielding vector pETDuet-1-PKRFP/EcoR. With pETDuet-1-PKRFP expression vector as template and primers of HS-F and HS-R, PCR fragment was obtained and subsequently digested with *Eco*R I and *Xho* I and ligated into pETDuet-1-PKRFP/EcoR yielding pETDuet-1-PKRFP-HF which contains one stem loop (5’-GCCGCTGTAGCTCTAGAGACAGCGGC-3’) after E*co*R I site (suppl data2).

RNA binding proteins (RBPs) are mainly composed of RNA binding scaffold (RBS) and auxiliary domain. RBS has a high degree of structural modularity, containing one or more RNA binding domains (RBD), such as the Pumilio/fem-3 binding factor (PUF)(Glisovic et al. 2008). The modular RBD of PUF is base specific, and amino acids at positions 12 and 16 in each repeat structure corresponds to different bases, for example, cysteine and glutamine corresponding to adenine, asparagine and glutamine corresponding to uracil, serine and glutamic acid corresponding to guanine, serine and arginine corresponding to cytosine (Dong et al. 2011; Filipovska et al. 2011). The coding sequence of engineered RNA binding scaffold1 (ERBS1) targeting RNA sequence of 5’-ACAGCGGC-3’ preceding to the A/U-rich region and internal SD, was designed using human PUM1 (pdb: 1M8Y) as backbone (Dong et al. 2011; Filipovska et al. 2011), and synthesized (suppl data3). The synthetic fragment was incorporated into pETDuet-1 vector with *Bam*H I and *Hin*d III double digestion and ligation, yielding pETDuet-1-ERBS1 expression vector.

PKRFP sequence was obtained from pETDuet-1-PKRFP by *Nde* I and *Xho* I double digestion, and further ligated into the second cloning site of pETDuet-1-ERBS1. This ERBS1 and PKRFP co-harboring expression vector was designated as pETDuet-1-ERBS1-PKRFP.

### Cell growth and protein expression

Protein expression was carried out as described previously (Sun and Leyh 2006). *E. coli* Rosetta (DE3) harboring expression vector was cultured in LB medium supplemented with 0.1 mg/mL ampicillin overnight at 37°C. After the culture OD_600_ reached 0.8, isopropyl-β-D-1-thiogalactopyranoside (IPTG) was added to a final concentration of 0.5 mmoles/L to induce protein expression for 3 h. Cells were pelleted by centrifugation (5,000 g) at 4°C for 10 min. Cell pellets were re-suspended in lysis buffer (50 mmoles/L potassium phosphate, 0.4 moles/L potassium chloride, 290 mmoles/L PMSF, 1.5 mmoles/L pepstatin A, 0.1 mg/mL lysozyme, pH 8.0) and incubated at 4°C for 1 h, and further disrupted by sonication (450 W,10 min) at 0°C. Cell debris was removed by centrifugation (18,000 g) at 4°C for 40 min. The supernatant was analyzed for RFP fluorescence.

### Nucleic Acid Thermal Stability Analysis

Two synthesized oligoes (HS1: 5’-GCCGCTGT-3’ and HS2: 5’-ACAGCGGC-3’) were mixed with equal concentrations, and absorbance at 260nm were recorded (Cary400, Agilent) upon temperature increasing from 25 to 90°C. Data were processed with equation (Souza et al. 2016): Fd+(Fn-Fd)/(1+exp((T-Tm)/a)). Where Fd is the denatured double strand reading at 260 nm at 90°C, Fn is the undenatured double strand reading at 260 nm at 25°C, T is the real time temperature, Tm is the melting temperature, and a is curvature parameter.

### RFP Fluorescence analysis

The supernatant from IPTG induced *E. coli* cells after was taken for RFP analysis using fluorescence spectrophotometer (Cary Eclipse, Agilent) with excitation wavelength of 550 nm, emission wavelength of 570-700 nm at 25°C.

After dripping bacterial suspension into the center of the clean slide, covering the cover slide and placing upside down on the stage of laser scanning confocal microscopy (LSM880, Zeiss, Germany), RFP expression in cells by microscopy was analyzed and cell images were recorded (excitation wavelengths of 561nm and emission wavelengths of 582-754nm).

### Protein SDS-PAGE analysis

One milliliter *E. coli* culture sample was taken after 3 h IPTG induction, and cells were pelleted by centrifugation (5,000 g). Cells were re-suspended with 100 μL 50 mmoles/L potassium phosphate buffer (pH 8.0), and protein loading buffer was added to mix well. Samples were denatured at 100 °C for 10 minutes, and centrifuged at 12000 g for 3 minutes. Supernatant (15 μL) and protein marker were loaded to the gel wells for electrophoresis. After electrophoresis, gel was carefully removed and stained with coomassie brilliant blue R-250. Gel photos were taken after washed with water and destained with de-staining solution.

## Results and discussion

### Rationale design of the experiment

Proteins, one of the major macromolecules, are essential for life. Protein translation could be regulated at different stages, initiation, elongation, termination, and post translational modification. Internal SD had been proposed and found play a role in initiating the downstream gene (Diwan and Agashe 2016; Cicek et al. 2013; Leith et al. 2019). To further characterize the upstream elements effects on the translation initiation role of internal SD, following vectors were constructed.

Firstly, PK coding sequence from *E. coli* was fused to RFP coding sequence (Fig.1a, suppl data1). One fragment, containing translation initiation needed mRNA elements which include an A/U-rich region, SD sequence and the start codon AUG, was introduced upstream of RFP coding sequence. A stop codon UGA resulting PK translation termination at 65nt of RFP coding sequence was used to avoid RFP fluorescence fusion protein formation which might confuse the ISTIR analysis (Fig.1a and d). Based on this design, RFP signal is positively correlated to ISTIR. In another words, RFP signal should be observed if internal SD can initiate the translation of downstream gene.

**Fig. 1.**
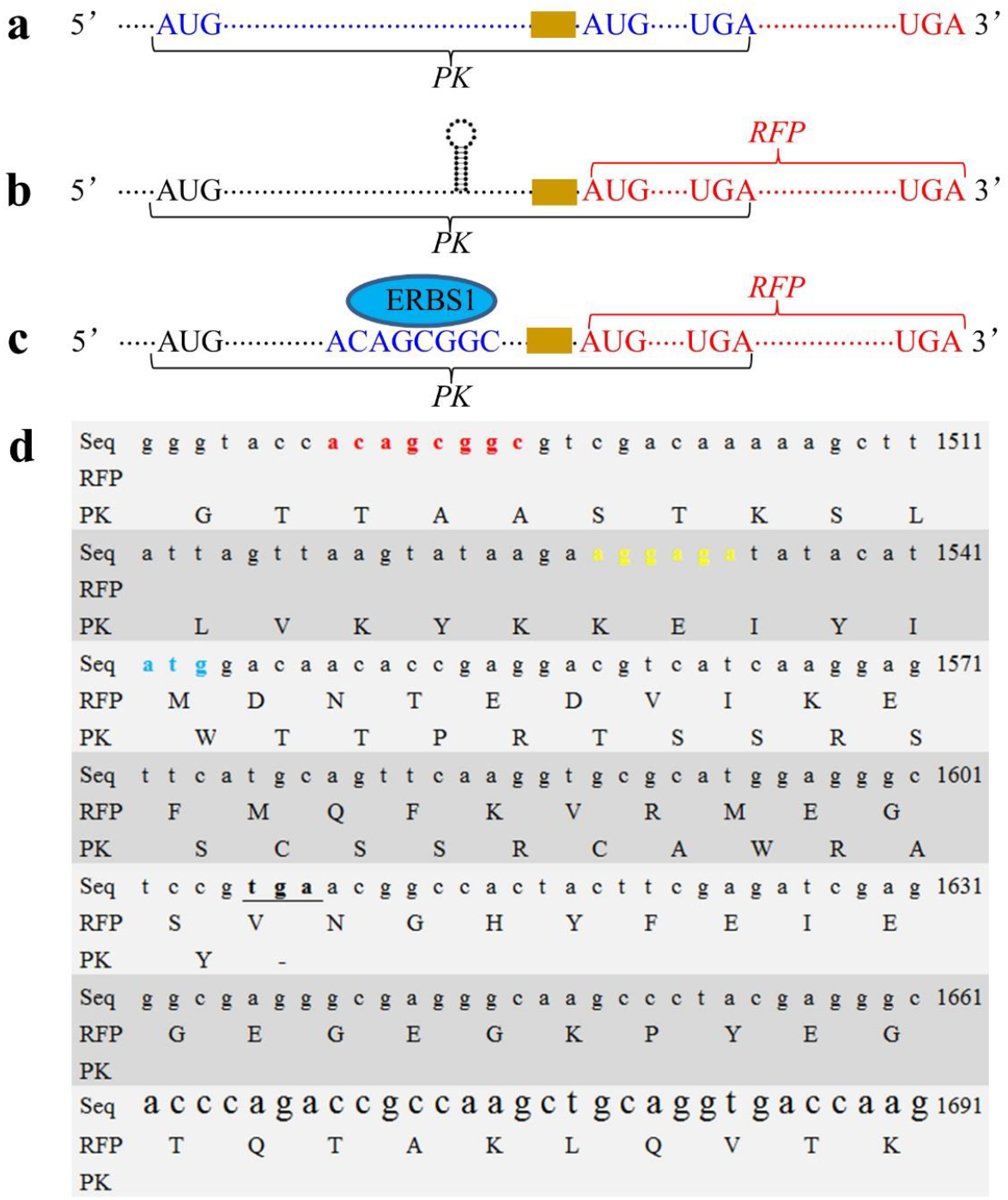
Illustration of fusion PKRFP sequence. a. Sequence in pETDuet-1-PKRFP, Brown box represents an AU-rich region and internal SD sequence; b. Sequence in pETDuet-1-PKRFP-HS with stem loop structure introduced; c. Sequence in pETDuet-ERBS-1-PKRFP; d. Detailed nucleotide sequence in the ERBS1 binding site, SD site, start codon of RFP and termination codon of PK. Numbers on the right denote the nucleotide position in PKRFP nucleotide sequence. Seq, the nucleotide sequence; RFP, RFP amino acid sequence; PK, PK amino acid sequence. RFP start codon was colored as light blue, internal SD was colored as yellow, ERBS1 binding site was colored as red. PK stop codon was underlined and bold.

Secondly, to assay the secondary structure effects on the ISTIR, one stem loop sequence was inserted into the upstream of the A/U-rich region and internal SD. Further IPTG induction was used to study functions of this secondary structure on ISTIR (Fig.1b, suppl data2).

Thirdly, ERBS1 (suppl data3) was designed to interact fragment of 5’-ACAGCGGC-3’ located at the upstream of the A/U-rich region and internal SD, and co-expressed with *PKRFP* to assay the ERBS1 impediment effects on the ISTIR (Fig.1c).

### Internal SD does not initiate the downstream RNA translation

*E. coli* cells harboring pETDuet-1-PKRFP expression vector was cultured, induced and analyzed for RFP expression, and it was shown that these cells could express PK upon IPTG induction, but no RFP protein was detected by SDS-PAGE (Fig.2, lane 1 and 2). These results indicated that even though the translation factors, such as the A/U-rich region, SD sequence and the start codon AUG were all present, the internal SD here could not initiate downstream gene translation. These surprising results are inconsistent with previous published results (Diwan and Agashe 2016; Cicek et al. 2013; Leith et al. 2019).

**Fig. 2.**
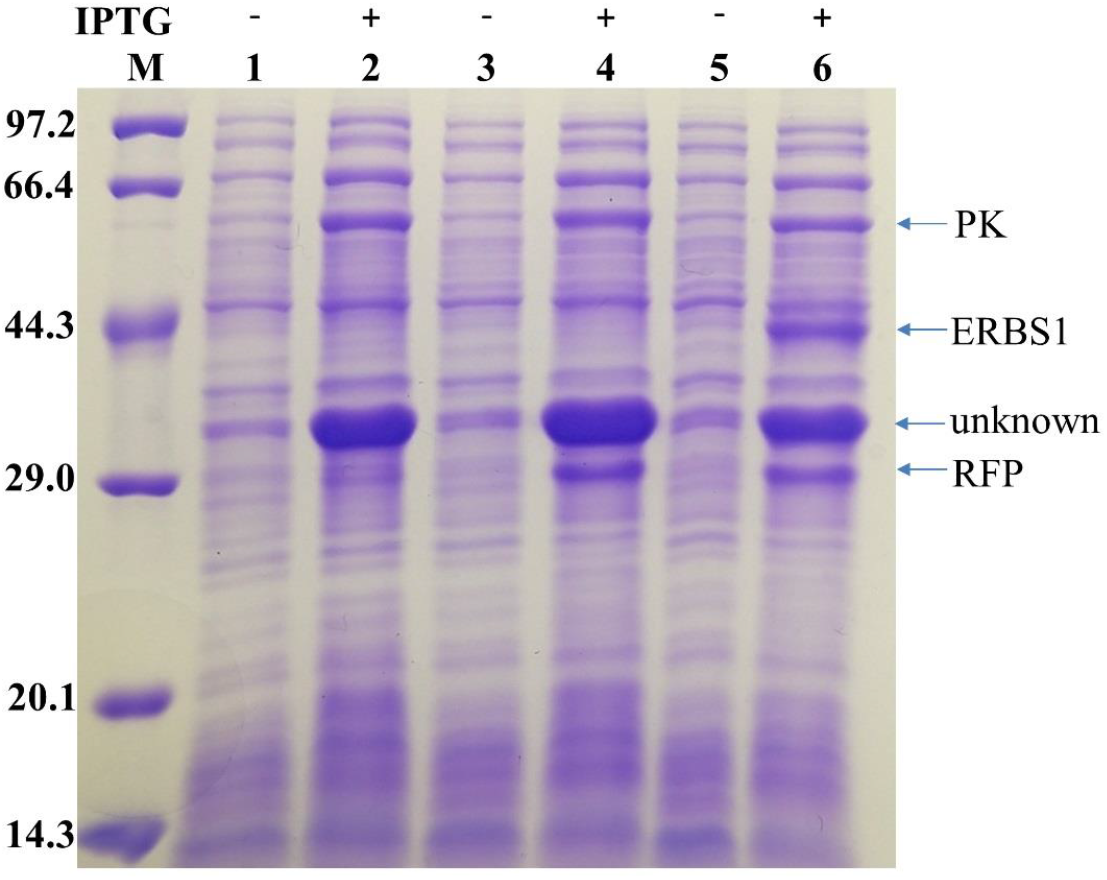
Protein expression with or without IPTG induction. Numbers on the left are protein marker size (kilodalton). Lane M, protein marker; lane 1, pETDuet-1-PKRFP without IPTG induction; lane 2, pETDuet-1-PKRFP with IPTG induction; lane 3, pETDuet-1-PKRFP-HS without IPTG induction; lane 4, pETDuet-1-PKRFP-HS with IPTG induction; lane 5, pETDuet-ERBS1-PKRFP without IPTG induction; lane 6, pETDuet-ERBS1-PKRFP with IPTG induction.

We also found that when PK translation was induced, one unknown protein was overexpressed (Fig.2, lane 1 and 2) which need to be further analyzed for its information, what this protein is and why it appeared with IPTG induction.

### Upstream stem loop activates internal SD downstream RNA translation

In order to assay if the internal SD presented in the coding sequence works properly or not, an RNA stem loop was introduced upstream of the A/U-rich region and internal SD. The rationale is that if it works fine, the downstream RFP should be translated, since the stem loop will diminish the possible interference from the upstream ribosome translation.

To test the designed complementary sequence will form stem loop or not, the oligoes were synthesized. Further Tm analysis showed the Tm of 57.90 (±1.90)°C, and the presence of absorbance jump (Fig.3a) indicated the break of the double strands which suggested the formation of stem loop. Cells harboring vector pETDuet-1-PKRFP-HS were cultured, induced by IPTG and assayed for RFP signal. As expected, in presence of stem loop structure, RFP was detected either by SDS-PAGE (Fig.2, lane 4), fluorescence detected by fluorometer (Fig.3b) and by confocal (Fig.3d). While these RFP signal were absent in the control *E. coli* cells (Fig.3b and 3c), and this is consistent with the SDS-PAGE result of Fig.2(lane 2). These results suggested that the internal SD designed here works. Internal SD induced internal translation initiation may result from ribosome translation pause preceding to internal SD.

**Fig. 3.**
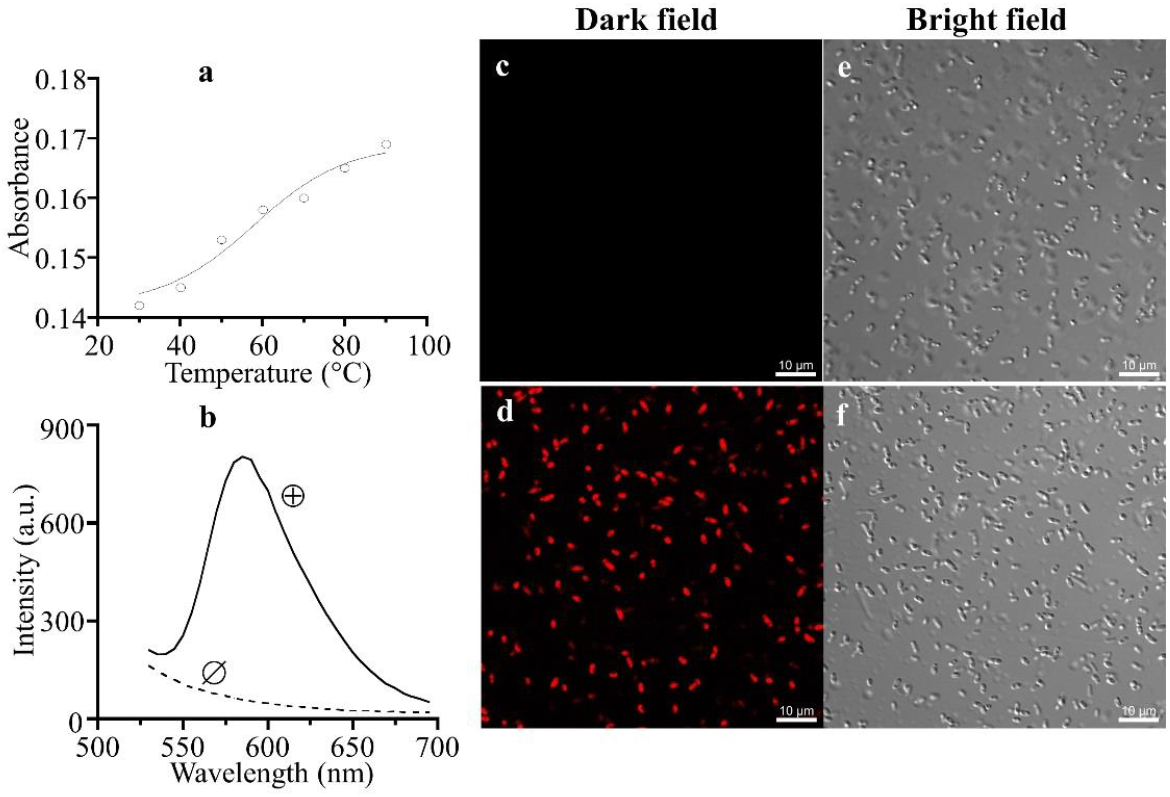
RFP fluorescence analysis in presence or absence of stem loop structure. a. Tm analysis with complementary oligoes. b. Fluorometer analysis on supernatant from cells harboring pETDuet-1-PKRFP (Ø) or pETDuet-1-PKRFP-HS (⊕) after induction by IPTG. Ex: 550 nm, Em: 570-700 nm. c and e, visualization of cells harboring pETDuet-1-PKRFP after induction with IPTG. d and f, visualization of cells harboring pETDuet-1-PKRFP-HS after induction with IPTG. Objective magnification: 100×, Numerical Aperture: 1.45.

### ERBS1 expression induced internal SD downstream RNA translation

Based on the stem loop blocking experiment, it seems that efficient upstream translation would deactivate the ISTIR. Impediments for ribosomes during translation will induce ribosome stalling on the mRNA instead of continuing protein synthesis (Muller et al. 2021). To assay if binding of RNA binding scaffold to the upstream of the A/U-rich region and internal SD would activate the ISTIR or not, ERBS1 recognizing the sequence of 5’-ACAGCGGC-3’ which located -53 of RFP start codon, was designed based on the human PUM1 structure and its coding sequence was synthesized. ERBS1 was subsequently ligated into the first multiple cloning site of pETDuet-1-PKRFP, constructing the co-expression vector pETDuet-ERBS1-PKRFP (Fig.1c).

Cells harboring vector pETDuet-ERBS1-PKRFP were cultured, induced by IPTG and assayed for protein expression. ERBS1, PK and RFP were all successfully expressed upon IPTG induction (Fig.2, lane 6). RFP expression was detected by fluorometer (Fig.4a) and confocal (Fig.4b and c). No RFP signal was detected for cells harboring pETDuet-1-PKRFP after IPTG induction (Fig.4a and b), which is consistent with the results of SDS-PAGE (Fig.2, lane 2). RFP signal was detected in cells harboring pETDuet-ERBS1-PKRFP after IPTG induction (Fig.4a and c), which is consistent with results of SDS-PAGE (Fig.2, lane 6). These results clearly demonstrated that the ERBS1 was functionally expressed and binds to the target sequence upstream of start codon of RFP, which activates ISTIR as the stem loop does shown above (Fig.2, lane 4; Fig.3b and 3c).

**Fig. 4.**
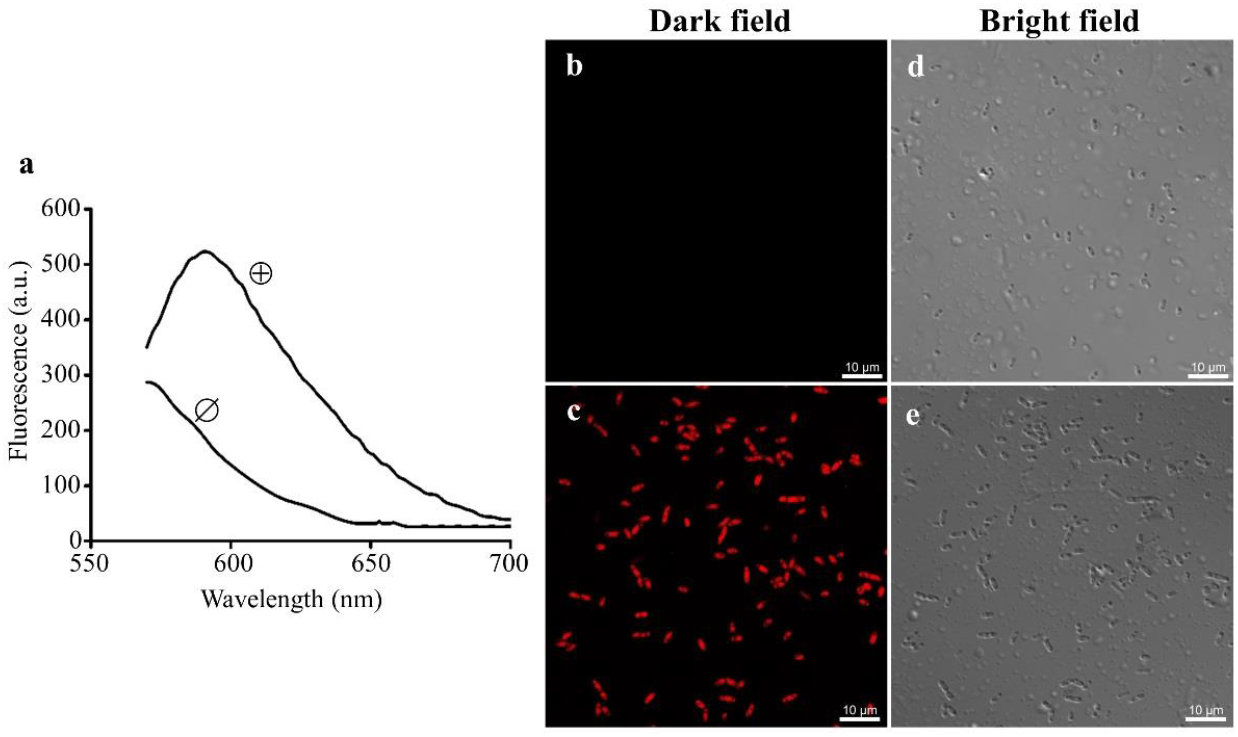
RFP fluorescence analysis in the presence or absence of cellular ERBS1. a. Fluorometer analysis on supernatant from cells harboring pETDuet-1-PKRFP (Ø) or pETDuet-ERBS1-PKRFP (⊕) after induction by IPTG (Ex: 550 nm, Em: 570-700 nm). b and d, visualization of cells harboring pETDuet-1-PKRFP after induction with IPTG. c and e, visualization of cells harboring pETDuet-ERBS1-PKRFP after induction with IPTG. Objective magnification: 100×, Numerical Aperture: 1.45.

Many methods had been developed to analyze the interactions between RBPs and their potential RNA targets (Ramanathan et al. 2019). The result that ERBS1 activates ISTIR (Fig.2, lane6 and Fig.4a and 4c) is interesting, and it may provide new strategy for testing interactions between RBPs and their target RNA molecules *in vivo*.

## Acknowledgement

This project was supported by Natural Science Foundation of Zhejiang province (grant no. LY19C050001).

