## Supplementary material for "Upstream ribosome impediments activate roles of internal Shine-Dalgarno sequence for translation initiation in *E. coli*": suppl table1

Supplementary table1

Primers used in this study

| Primer name | Primer sequence(5’-3’) |
| --- | --- |
| PK-F | GGAATTCCATATGATGTCCAGAAGGCTTCGCAGAAC |
| PK-R | CGGGGTACCCTCTACCGTTAAAATACGCGTG |
| RFP-F1 | CGGGGTACCACAGCGGCGTCGACAAAAAGCTTATTAGTTAAGTATAAGAAGGAG |
| RFP-F2 | ATTAGTTAAGTATAAGAAGGAGATATACATATGGACAACACCGAGGACGTC |
| RFP-R | CCGCTCGAGCTGGGAGCCGGAGTGGCGGGCC |
| ER1 | GTATTTTAACGGTAGAGGAATTCGGTACCACAGCGGCGTC |
| ER2 | GACGCCGCTGTGGTACCGAATTCCTCTACCGTTAAAATAC |
| HS-F | GAATTCGCCGCTGTAGCTCTAGAGACAGCGGCGTCGACAAAAAG |
| HS-R | CTCGAGCTGGGAGCCGGAGTGGC |
