## Supplementary material for "Upstream ribosome impediments activate roles of internal Shine-Dalgarno sequence for translation initiation in *E. coli*": suppl data1

Supplementary data1

Sequence of PKRFP

CATATGTCCAGAAGGCTTCGCAGAACAAAAATCGTTACCACGTTAGGCCCAGCAACAGATCGCGATAATAATCTTGAAAAAGTTATCGCGGCGGGTGCCAACGTTGTACGTATGAACTTTTCTCACGGCTCGCCTGAAGATCACAAAATGCGCGCGGATAAAGTTCGTGAGATTGCCGCAAAACTGGGGCGTCATGTGGCTATTCTGGGTGACCTCCAGGGGCCCAAAATCCGTGTATCCACCTTTAAAGAAGGCAAAGTTTTCCTCAATATTGGGGATAAATTCCTGCTCGACGCCAACCTGGGTAAAGGTGAAGGCGACAAAGAAAAAGTCGGTATCGACTACAAAGGCCTGCCTGCTGACGTCGTGCCTGGTGACATCCTGCTGCTGGACGATGGTCGCGTCCAGTTAAAAGTACTGGAAGTTCAGGGCATGAAAGTGTTCACCGAAGTCACCGTCGGTGGTCCCCTCTCCAACAATAAAGGTATCAACAAACTTGGCGGCGGTTTGTCGGCTGAAGCGCTGACCGAAAAAGACAAAGCAGACATTAAGACTGCGGCGTTGATTGGCGTAGATTACCTGGCTGTCTCCTTCCCACGCTGTGGCGAAGATCTGAACTATGCCCGTCGCCTGGCACGCGATGCAGGATGTGATGCGAAAATTGTTGCCAAGGTTGAACGTGCGGAAGCCGTTTGCAGCCAGGATGCAATGGATGACATCATCCTCGCCTCTGACGTGGTAATGGTTGCACGTGGCGACCTCGGTGTGGAAATTGGCGACCCGGAACTGGTCGGCATTCAGAAAGCGTTGATCCGTCGTGCGCGTCAGCTAAACCGAGCGGTAATCACGCAAACCCAGATGATGGAGTCAATGATTACTAACCCGATGCCGACGCGTGCAGAAGTCATGGACGTAGCAAACGCCGTTCTGGATGGTACTGACGCTGTGATGCTGTCTGCAGAAACTGCCGCTGGGCAGTATCCGTCAGAAACCGTTGCAGCCATGGCGCGCGTTTGCCTGGGTGCGGAAAAAATCCCGAGCATCAACGTTTCTAAACACCGTCTGGACGTTCAGTTCGACAATGTGGAAGAAGCTATTGCCATGTCAGCAATGTACGCAGCTAACCACCTGAAAGGCGTTACGGCGATCATCACCATGACCGAATCGGGTCGTACCGCGCTGATGACCTCCCGTATCAGCTCTGGTCTGCCAATTTTCGCCATGTCGCGCCATGAACGTACGCTGAACCTGACTGCTCTCTATCGTGGCGTTACGCCGGTGCACTTTGATAGCGCTAATGACGGCGTAGCAGCTGCCAGCGAAGCGGTTAATCTGCTGCGCGATAAAGGTTACTTGATGTCTGGTGACCTGGTGATTGTCACCCAGGGCGACGTGATGAGTACCGTGGGTTCTACTAATACCACGCGTATTTTAACGGTAGAGGGTACCACAGCGGCGTCGACAAAAAGCTTATTAGTTAAGTATAAGAAGGAGATATACATATGGACAACACCGAGGACGTCATCAAGGAGTTCATGCAGTTCAAGGTGCGCATGGAGGGCTCCGTGAACGGCCACTACTTCGAGATCGAGGGCGAGGGCGAGGGCAAGCCCTACGAGGGCACCCAGACCGCCAAGCTGCAGGTGACCAAGGGCGGCCCCCTGCCCTTCGCCTGGGACATCCTGTCCCCCCAGTTCCAGTACGGCTCCAAGGCCTACGTGAAGCACCCCGCCGACATCCCCGACTACATGAAGCTGTCCTTCCCCGAGGGCTTCACCTGGGAGCGCTCCATGAACTTCGAGGACGGCGGCGTGGTGGAGGTGCAGCAGGACTCCTCCCTGCAGGACGGCACCTTCATCTACAAGGTGAAGTTCAAGGGCGTGAACTTCCCCGCCGACGGCCCCGTAATGCAGAAGAAGACTGCCGGCTGGGAGCCCTCCACCGAGAAGCTGTACCCCCAGGACGGCGTGCTGAAGGGCGAGATCTCCCACGCCCTGAAGCTGAAGGACGGCGGCCACTACACCTGCGACTTCAAGACCGTGTACAAGGCCAAGAAGCCCGTGCAGCTGCCCGGCAACCACTACGTGGACTCCAAGCTGGACATCACCAACCACAACGAGGACTACACCGTGGTGGAGCAGTACGAGCACGCCGAGGCCCGCCACTCCGGCTCCCAGCTCGAG
