## Supplementary material for "Upstream ribosome impediments activate roles of internal Shine-Dalgarno sequence for translation initiation in *E. coli*": suppl data3

Supplementary data3

Sequence of ERBS1

CATATGGGTCGTAGCCGCCTGCTGGAAGATTTTCGCAACAACCGCTACCCGAACCTGCAGCTGCGCGAAATTGCGGGTCATATTATGGAGTTTAGCCAGGATCAATATGGCTGCTATGTGATTCGCCATGTGCTGGAACGCGCGACCCCGGCGGAACGCCAGCTGGTGTTTAACGAAATTCTGCAGGCGGCGTATCAGCTGATGGTGGATAAATTCGCGAGCAACGTGGTGGAAAAATGCGTGGAATTCGGCAGCCTGGAACAGAAACTGGCGCTGGCAGAACGTATTCGTGGCCATGTGCTGTCTCTGGCACTGCAGAAATTCGCATCTAACGTGGTTGAAAAATGTGTTGAATTCATTCCGTCTGATCAGCAGAACGAAATGGTTCGTGAACTGGATGGCCATGTTCTGAAATGCGTTAAAGATCAGTATGGCTGCTACGTTATTCGTCATGTTCTGGAATGCGTTCAGCCGCAGTCTCTGCAGTTCATCATCGATGCATTCAAAGGCCAGGTTTTCGCACTGTCTACCCACAAATTCGCCTCCAACGTTGTTGAAAAATGCGTTGAACACTGCCTGCCGGACCAGACCCTGCCGATCCTGGAAGAACTGCACCAGCACACCGAACAGCTGGTTCAGGACATGTATGGCTGTCGCGTTATCCAGAAAGCCCTGGAACACGGTCGTCCGGAGGACAAATCCAAAATCGTAGCCGAGATCCGTGGTAACGTACTGGTACTGTCCCAACACCAATACGGTTGTTACGTAATCCGTCACGTACTGACCCACGCTTCCCGTACCGAGCGTGCTGTACTGATCGACGAGGTATGTACGATGAATGACGGTCCGCACTCCGCTCTGTACACTATGATGAAAGACCCTTACGGTTGTCGTGTCATCCAACGTATCCTGGACGTCGCTGAGCCAGGTCAACGTAAAATCGTCATGCACAAAATCCGTCCACACATCGCTACTCTGCGTAAATACACTTACGGCAAGCACATCCTGGCTAAGCTGGAGAAGTACTACATGAAGAATGGTGTCGACCTGGGTAAGCTT
